# Blended Bioinformatics Training in Resource-Limited Settings: A case study of challenges and Opportunities for Implementation

**DOI:** 10.1101/431361

**Authors:** Azza Ahmed, Ayah A. Awadallah, Mawada T. Elmahdi, Maram A. Suliman, Atheer E. Khalil, Hassan Elsafi, Basil D. Hamdelnile, Mohamed Abdullateif, Faisal M. Fadlelmola

## Abstract

**Motivation:** Delivering high quality distance-based courses in resource limited settings is a challenging task. Besides the needed infrastructure and expertise, effective delivery of a bioinformatics course could benefit from hands-on sessions, interactivity, and problem-based learning approaches.

**Results:** In this article, we discuss the challenges and best practices in delivering bioinformatics training in limited resource settings taking the example of hosting and running a multiple-delivery online course, Introduction to Bioinformatics (IBT), that was developed by the H3ABioNet Education and Training Working Group and delivered in 27 remote classrooms across Africa in 2017. We take the case of the University of Khartoum classroom. Believing that our local setting is similar to others in less developed countries, so we also reflect upon aspects like classroom environment and recruitment of students to maximize outcomes.

**Supplementary information:** Supplementary data are available

## 1 Introduction

The rapid advancements in genomics and molecular biology research and applications necessitates adequate, up-to-date and complimentary training in biology and computer science [Attwood *et al.*, 2017, Mulder *et al.*, 2018].

Physical face-to-face bioinformatics training workshops are one way to address this need, by providing opportunities for networking and first hand discussions[Brazas and Ouellette, 2013]. However, when run in settings of limited access to local bioinformatics expertise, funding and proper infrastructure [Bishop *et al.*, 2014], this model becomes very expensive to run and modest in students’ intake. Of question is the relevancy and applicability of skills acquired from such a training to the bioinformatician’s own environment in the long-term [Gurwitz *et al.*, 2017].

Learning methods connecting distant learners and educators have evolved with technology, from postal services to Massively Open Online Courses (MOOCs) [Moore and Kearsley, 2011]. For bioinformatics training, both edX and Coursera, 2 popular MOOC providers, offer complete specializations for both biologists (https://www.edx.org/micromasters/bioinformatics) and computer scientists (https://www.coursera.org/specializations/bioinformatics). Yet, some MOOCs may be inaccessible for developing countries’ learners due to conditions like technological access, digital literacy, cultural relevance and social identity threats [Castillo *et al.*, 2015, Kizilcec *et al.*, 2017], and other typical caveats associated with MOOCs - competing priorities of learners and information overload [Hew and Cheung, 2014a].

H3ABioNet, the pan African Bioinformatics Network [Mulder *et al.*, 2015], is making strides towards bridging the bioinformatics training gap in Africa by designing and offering a 3-months multiple-delivery-mode training course, Introduction to Bioinformatics (IBT), across all its nodes including Sudan [Gurwitz *et al.*, 2017]. The IBT model blends local in-person tutoring sessions with online elements delivered via the course website (https://training.h3abionet.org/IBT_2017/), the on-line learning management system, Vula (http://www.cilt.uct.ac.za/cilt/vula), and the open source videoconferencing system, Mconf(https://mconf.sanren.ac.za/)

This study assesses the efficiency and effectiveness of the 2017 iteration of the IBT (IBT_2017) training model from the local learner’s and Teaching Assistants’ (TAs) perspectives in the H3ABioNet Node of Sudan based at the University of Khartoum. We investigated factors contributing to training success, as inferred from responses to surveys disseminated to both local learners and TAs. Our results agree with empirical data suggesting that local group discussions improved students’ ability to access the course materials [Yousef *et al.*, 2015], especially when facilitated by volunteering course alumni [Murugesan *et al.*, 2017]. Also, the African context ingrained into the course made the content relevant to the local learners, and hence the training aligned with their expectations [Castillo *et al.*, 2015]. Our setting resembles others in less developed countries, so we further reflect upon aspects like classroom environment and recruitment of learners to maximize outcomes.

## 2 Relevant literature

The effectiveness of MOOCs in less developed countries is hindered by barriers of technology and context[Castillo *et al.*, 2015]. Efforts addressing these barriers include +Acumen (https://www.plusacumen.org), which aims at empowering social change and provides MOOCs employing in-video transcripts, culturally-diverse case studies, and content that is viewable off-line and platform-agnostic. Consequently, +Acumen attracts participants from a diverse pool of countries, including Afghanistan, Botswana and Sri Lanka [Cheney, 2017]. Successful participation from those countries and other top Fragile States [OECD, ????] was also reported in AuthorAID’s offering on Scientific research writing, which utilized low-bandwidth friendly format via mainly text-based content, and voluntary course alumni as facilitators [Murugesan *et al.*, 2017].

Interestingly, Kizilcec *et al.* 2017 have demonstrated significant improvement in persistence and completion rates of less-developed countries’ learners in MOOCs by brief psychological interventions to lessen social identity threats, like value affirmations and social belonging [Kizilcec *et al.*, 2017]. Furthermore, studies reporting on the application of the blended MOOC (bMOOC) paradigm, combining online MOOC components with in-class interactions, have systematically shown positive educational indicators; even when applied in resource limited settings [Ghadiri *et al.*, 2013, Yousef *et al.*, 2015, Nkuyubwatsi, 2016].

Especially for bioinformatics, there are huge, urgent, unmet training needs[Attwood *et al.*, 2017] exacerbated by the breadth of the discipline and its rapid evolution [Mulder *et al.*, 2018], notwithstanding difficulty of curricula design to learners of diverse backgrounds [Bishop *et al.*, 2014], and the shortage in experienced qualified trainers [Attwood *et al.*, 2017]. Therefore, various international efforts were exerted to bridge this skills gap, like GOBLET, ELIXIR, BD2K TCC and H3ABioNet. These efforts focused on short face-to-face and online courses which is preferable by researchers in their later career stages [Attwood *et al.*, 2017].

However, For basic bioinformatics users [Welch *et al.*, 2014], both Coursera and edX provide introductory level training. Tables 1 and 2 compare these offerings with H3ABioNet’s bMOOC in different aspects. Their accessibility to learners in less developed countries is yet to be investigated.

**Table 1.**
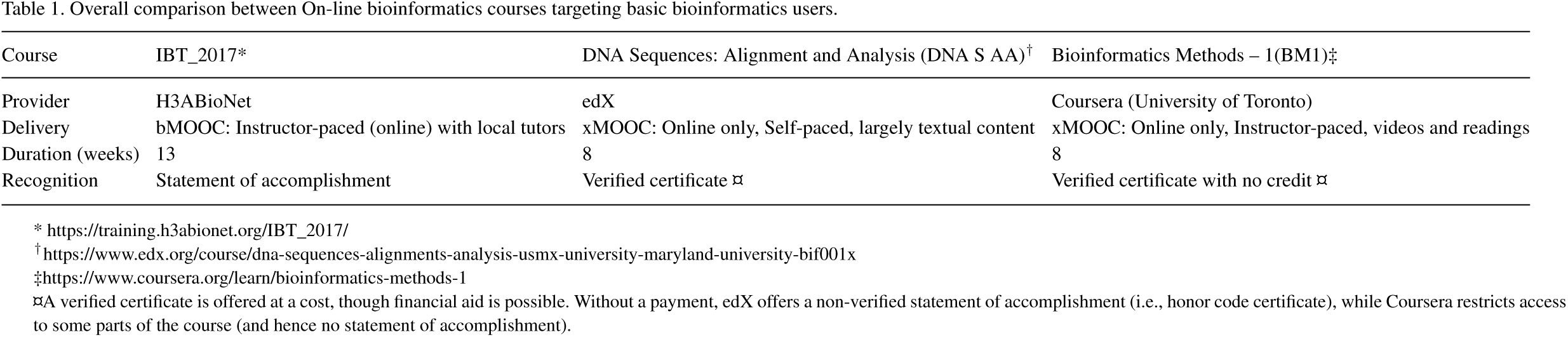
Overall comparison between On-line bioinformatics courses targeting basic bioinformatics users.

**Table 2.** Content-wise comparison between On-line bioinformatics courses targeting basic bioinformatics users

| Course | IBT_2017 | DNA S AA | BM1 |
| --- | --- | --- | --- |
| Genetics review | - | + | - |
| Databases | + | + | + |
| Linux | + | - | - |
| Sequence similarity | + | + | + |
| Genomics | + | + | + |
| Phylogenetics | + | + | + |
| Gene expression | Extra material | + | + |
| Protein structure | - | - | - |
| Selection analysis | - | - | + |
| Metagenomics | - | - | + |

- IBT\_2017: The *Introduction to Bioinformatics* course from H3ABioNet in its 2017 iteration.
- DNA S AA: The *DNA Sequences: Alignment and Analysis* course from edX.
- BM1: The *Bioinformatics Methods – 1* course from Coursera.

## 3 Methods

### 3.1 The 2017 iteration of the IBT

In its 2017 iteration, the IBT course started on May 9th, with 2 days/week for in-person interactive sessions. Building on its first iteration of 2016 [Gurwitz *et al.*, 2017], the IBT_2017 was composed of 6 modules: Introduction to databases and resources, Linux, Sequence alignment theory and application, Multiple sequence alignment, Genomics, Molecular evolution and phylogenetics (Table 2). The design, learning objectives and contents of these modules are already described in [Gurwitz *et al.*, 2017], so here we only comment on the local supporting set up of the classrooms.

In the H3ABioNet Node of Sudan, 73 students registered in the IBT_2017. The majority of these students (participants or learners herein) have been selected from a waiting list from the IBT 2016 iteration, based on their interest and basic understanding of the central dogma of molecular biology, and hence they came from diverse specializations (Figure 1 and Figure SF3), at different educational levels (6% were at the BSc level, 41% current MSc students, 11% current PhD students and 34% were MSc and PhD graduates not pursuing any degree) and career affiliations (66% in academic institutes, 4% in governmental ministries, 7% in Research centers, 3% in private companies and hospitals, with 12% unemployment rate) as shown in Figure 1 (C and A respectively). Figure 1B further shows that the highest educational institute for the majority of participants is the University of Khartoum (73%, compared to 13% from the other Sudanese universities, and 4% who studied abroad).

**Fig. 1.**
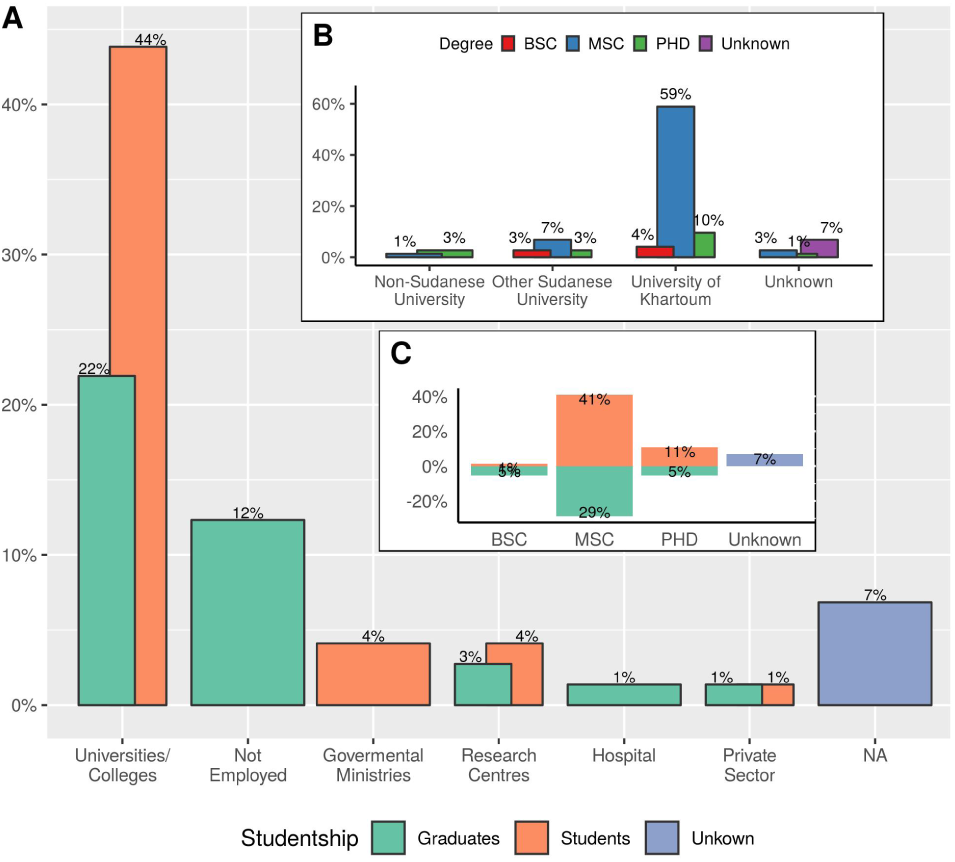
Demographics of the IBT_2017 participants in the H3ABioNet University of Khartoum Node, Sudan. A) Different affiliations of IBT participants stratified by their status as Current students or Graduates. Grouping into the shown categories was done by manual assignment of data to the appropriate category. B) The distribution of the institutes awarding the highest degree for the course participants. C) The distribution of the highest academic degree of the IBT participants stratified by their status as Current students or Graduates.

To accommodate this large number, two classrooms were set up for physical interactions within University of Khartoum main campus: the CBSB laboratory, equipped with 20 PCs and network ports to accommodate an additional 12 PCs/laptops; and the Main Library computer lab that can accommodate up to 70 participants. Collectively, these two locations hosted the 73 registered participants, split to 33 and 40 respectively. These two locations vary in their infrastructure as well: the PCs in the CBSB lab are appropriate for bioinformatics training and research with larger screen sizes and more CPU and memory capacities, while those in the Main library needed more effort from the local IBT team and University of Khartoum Information Technology Network Administration (ITNA) staff to set them up.

This larger intake (compared with 22 participants in 2016) has been managed by a local staff, which besides the node Principle Investigator, was composed of 7 teaching assistants (TAs) who are among the alumni of the IBT_2016 iteration with previous excellent background in genetics & molecular biology. Their prior IBT experience helped them provide actionable support to the new course participants, and their facilitation job was tremendously eased with the on-line staff training sessions provided by the IBT core team. Only a single system administrator was available for the duration of the IBT_2017, given the physical proximity of both classrooms.

### 3.2 Measurement and evaluation

While we lack data on the performance details of our IBT_2017 course participants (individual assessments and tests’ results were managed by the IBT core team in South Africa, and only shared directly with each participant), we have alternative aggregate data from the following sources:

1. The waiting list from the previous IBT_2016, containing demographic data on our course participants.
2. Three surveys designed to monitor our learners experience throughout the course. Those surveys were distributed at the start of the course (Supplementary SM1), the middle (Supplementary SM2) and end points (Supplementary SM3). Figure 2 tracks this data against participants’ attendance and withdrawal patterns (Supplementary SM7). Absence pattern is the difference between the total number of registered participants (73), and the present and withdrawn participants in each session, hence it is not explicitly shown.
3. There is also data identifying participants who earned a certificate upon successfully satisfying the course requirements.
4. An exit survey was also designed and disseminated to withdrawn course participants to capture the reasons motivating their withdrawal (Supplementary SM6).
5. Another survey was designed and disseminated to the local TAs, assessing the extent to which the course experience was valuable to them (Supplementary SM4).
6. Finally, a follow up survey was sent to the course participants 9 months upon their completion of the IBT_2017 (Supplementary SM5).

**Fig. 2.**
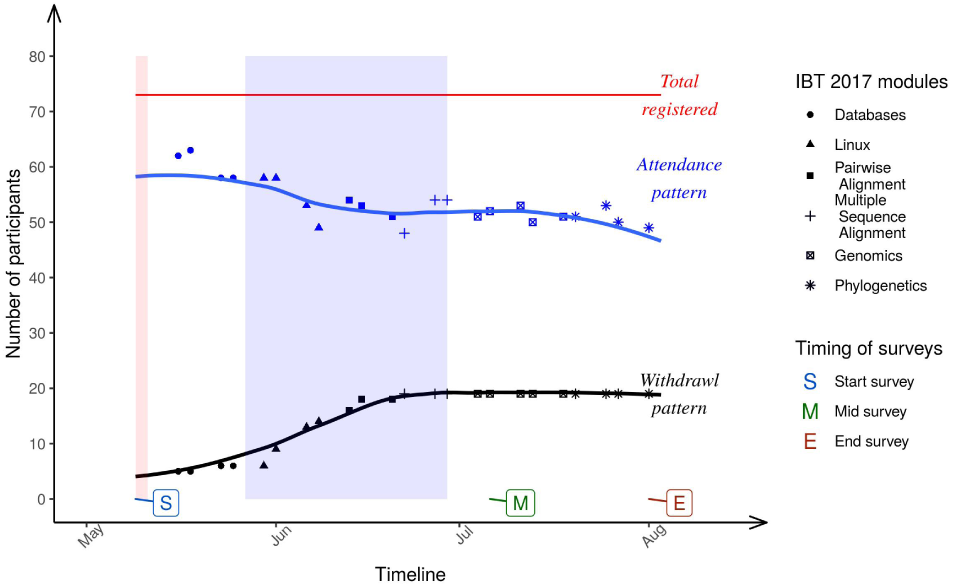
Patterns of attendance and withdrawals in the 6 IBT modules. Points of data collection (surveys) are also highlighted in the timeline. The red shaded area coincides with the orientation week, whereas the blue shaded area is the holy month of Ramadan.

Collectively, we used this data to investigate factors associated with a successful experience (or alternatively, failure to satisfy the course requirements or complete withdrawal), so that we are better informed for future course runs, or similar training initiatives.

## 4 Results

### 4.1 Attendance & Withdrawal patterns

Students attendance and retention is a major concern contributing to a successful MOOC experience [Hew and Cheung, 2014b] especially that to many learners, the problem is about committing the needed study hours despite their busy schedules.

In this 3 months course, we noted that for our local 73 registered participants, the attendance rate was higher at the beginning of the course (~85%), then it dropped progressively towards mid-June and early July (Figure 2), concurrent with gradual increase in withdrawal. Poor response rate from withdrawn students to the exit survey (Supplementary SM6) limits our ability to conclusively reason about it, except to note that it can be related to 2 factors: 1) Co-occurring with the Linux module, which requires a mode of thinking a bit alien to wet lab biologists. 2) Culturally, the IBT_2017 started just a few weeks before the holy month of Ramadan, coinciding with end-of-year/semester holidays in many universities and colleges. Taken in light of Figure 1A, during the few starting weeks it was easy to follow up with the progress of material and activities for students (53% of the total IBT_2017 participants)- both full-time (44%) and part time (9%), and other academic staff (22% of the total IBT_2017 participants). However, once these holidays were over, participants (collectively, 75% of the total) needed to be back to fulltime working hours and classes in their respective institutes, making it harder for them to attend IBT_2017 sessions in person and timely work towards their assessments and tests. Figure 2 shows that the withdrawal pattern plateaued after this point in time, totaling 5 and 14 participants from the CBSB and Main Library classrooms respectively, summing to 19 participants (26% of the total).

Remarkably, Figure 1B shows that the majority of the IBT_2017 participants (73%) have graduated from University of Khartoum, as do the local course staff. This could justify why ~63.6% of the participants heard of the course through friends, 27.3% from their supervisors or mentors, and the remaining 14.5% through social media (Supplementary SF1). It also suggests that local circles of friends/ acquaintances were already in place before the course had actually started. This support system in place could explain why for those participants who didn’t withdraw (54), only 3 participants (6%) failed the course requirements.

We investigate predictive models of learner’s performance (Success, Failure or Withdrawal) based on the demographics of the classrooms in the Discussion (Figure 5, Supplementary SF4, SF5, Mathematical models), noting poor response rate to the exit survey (Supplementary SM6) of only 1 response. We also note limitations impinged from unmoderated personal circumstances; like traveling, health problems and unwaivable work/study duties.

### 4.2 Participants’ perceptions & expectations

Here, data were collected from 3 surveys (Supplementary SM1, SM2, SM3) disseminated at the start of the course, the mid-point and the end (Figure 2). Out of the 73 participants, only 33 (45%) filled all the 3 surveys, while 15 (21%) never filled any (Supplementary SF2).

The experience of the IBT_2017 is both unique and new to our participants considering its blended multi-delivery learning model [Gurwitz *et al.*, 2017], and extended 3-months duration. Our surveys aimed to check the alignment between participants’ expectations and course scope [Via *et al.*, 2011]. On a labeled five-point Likert scale, we asked participants about their perspectives in terms of their prior experience level in each module (start survey); the extent to which the content was appropriate, and the level it met their expectations (in mid-course and end surveys, for each module taught until that point).

Participants’ perceptions on each of the six modules largely followed the same trend (Supplementary SF6 - SF11), hence their average across all modules is presented in Figure 3. Not surprisingly, most participants were initially largely unfamiliar with the various modules (with the 75th percentile of responses below neutral familiarity level), especially Linux (Supplementary SM8). Progressively however, we see higher satisfaction levels (with the median of the averaged responses at a level above Comfortable in Figure 3) in terms of appropriateness of the taught material and meeting participants expectations.

**Fig. 3.**
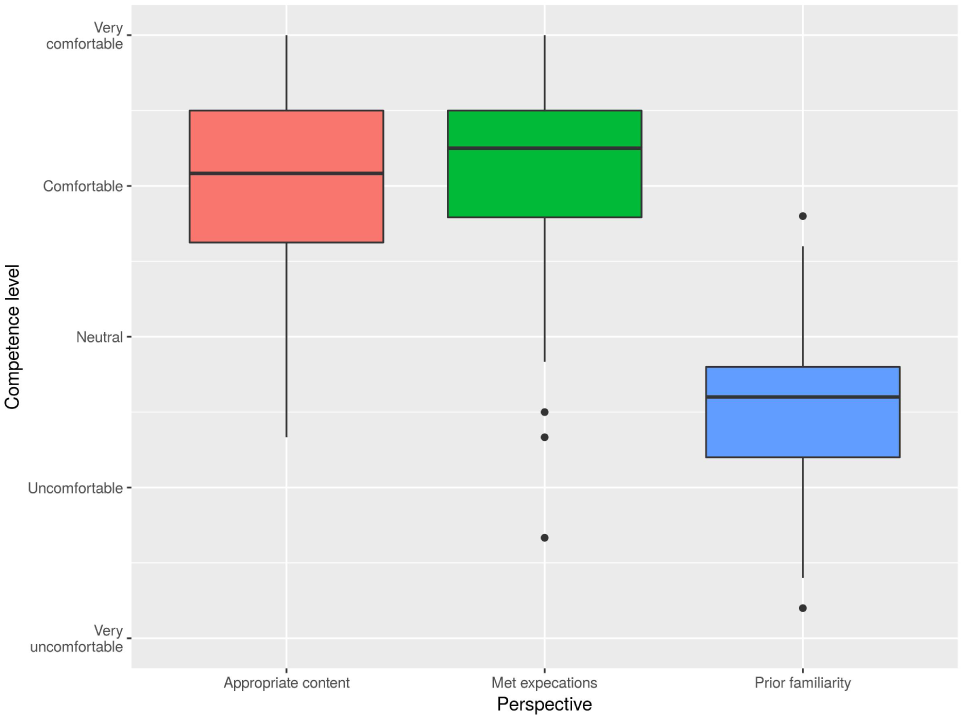
Average IBT participants’ perceptions on the 6 modules taught in the 2017 iteration of the course, in terms of their level of competence, measured as the level of their prior familiarity with the modules, and their satisfaction upon completion according to the responses collected via the Start, Mid-course and End surveys.

Another aspect, is the extent to which our participants made use of the local and remote classroom elements to ameliorate their learning experience. Namely, we were interested in: Networking with others and use of Vula. We monitored the progression in these aspects at the mid-course and end surveys. The responses depicted in Figure 4, show a positive trend as the course advanced, suggesting more familiarity with the blended MOOC model. Yet, we remark a modest and steady amount of networking with other IBT classrooms throughout the course (In total, 38 participants filled both the mid-course and end surveys, with 2 and 5 unique responses for each survey respectively (Supplementary SF2)).

**Fig. 4.**
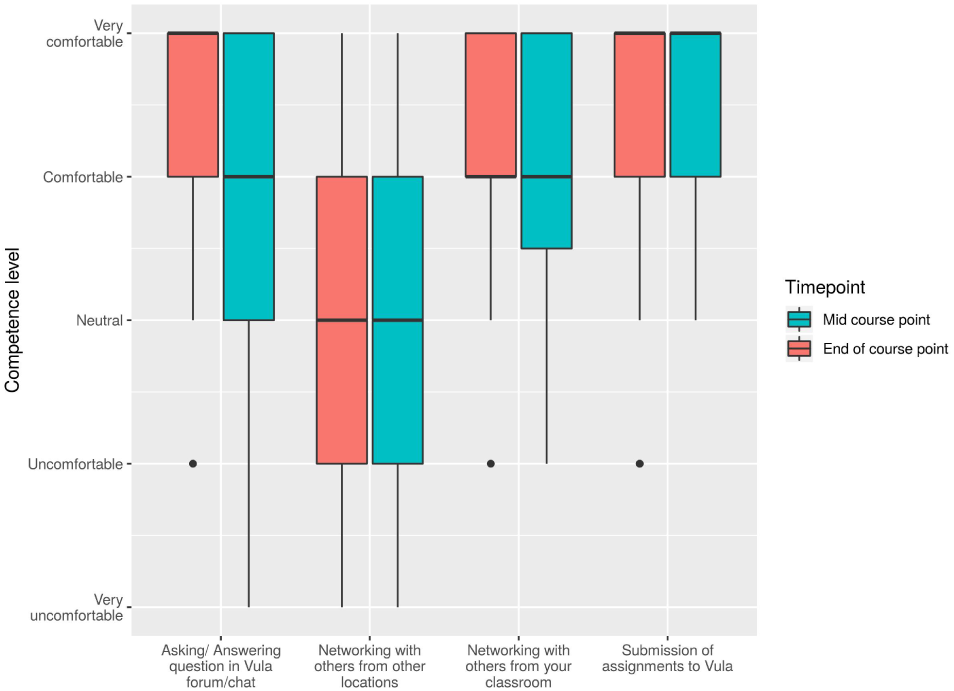
Students utilization levels of various elements of online learning: Vula and networking measurements were taken from surveys in the middle of the course and at the end of the course.

## 5 Discussion

### 5.1 Classroom demographics & performance

Consistent with the largely female students’ body in faculties related to natural sciences, health, and agriculture in Sudan [Huyer, 2015], the majority of the IBT_2017 participants in both classes of the H3ABioNet node of Sudan were females (80%); and they were also more likely to satisfy the course requirements in comparison with their male peers (Supplementary SF3). Utility of gender as a performance predictor can also be seen from Recursive Partitioning (rpart) tree [Therneau and Atkinson, 2018] of (Supplementary SF4, Listing1) built by selecting splitting covariates to minimize the Gini coefficient as an information measure.

However, when changing the partitioning algorithm to a conditional inference tree [Hothorn *et al.*, 2006], the location of the local IBT_2017 classroom (the CBSB lab or Main Library), is the most important covariate in predicting performance (Supplementary SF5). This is expected considering the inherent infrastructure differences between the 2 locations. The CBSB lab is designed to facilitate bioinformatics training and research in terms of stable internet connection and more powerful computers; whereas the Main Library classroom had Internet connectivity issues at the beginning of the course, which was frustrating to some of the participants (and in occasions encouraged some to withdraw early on).

Modeling differences of these algorithms (Supplementary SF4, SF5), are alterable to the high degree of class imbalance in the entire dataset (70% Success, 26% Withdrawal and 4% Failure), even when seen in each classroom independently (85%, 15% and 0% for CBSB class, and 58%, 35% and 8% for the Main Library respectively). We therefore built a multinomial classification model to further examine all demographic factors collectively (Figure 5, ST2 and Listing2). While a potential problem with this model is the assumptions of independence and constant performance, we see that both the physical location of the classroom and Gender are the only statistically significant demographic factors, with a note that MSc and PhD participants had higher odds of success than BSc level candidates; as did unemployed participants and those working in research centers or the private sector. Whether a participant is currently a student or has graduated from the said level had slight effect on their odds of success (see SF3).

**Fig. 5.**
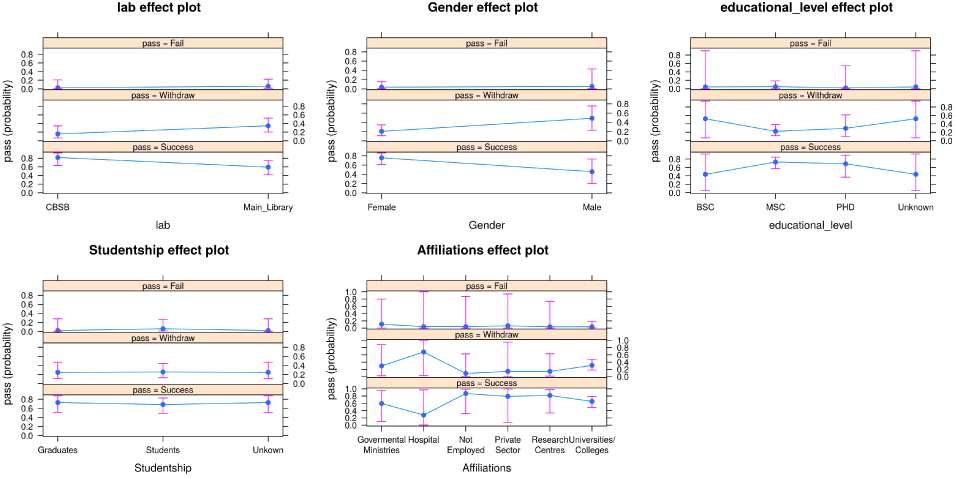
Effect of the logistics and demographics of the class (lab location, Gender, Educational level and Studentship) on IBT participants’ performance (Success, Withdrawal or Failure) based on the 10-fold cross validated multinomial model of performance of Supplementary Table ST1.

Figure 1A underscores that about half the entire local IBT_2017 classes (53%) are students, 15% of whom are part-timers with affiliations in either governmental ministries or research laboratories. We reckon the better odds of success for students to their interest in a specific problem at hand. We see no statistical evidence of interaction between part/full time status and studentship, hence it is not shown in the ST1 model. However, the ability of part-timers to satisfy the course requirements and acquire higher odds of success (Figure 5, supplementary SF3, SM8), suggest the appreciation of these institutes to staff training; and possibly hints at avenues for local sustainable research collaborations.

Yet, there is the 12% unemployment ratio among our participants. While we didn’t explicitly investigate the employability of typical biological sciences graduates in Sudan, by large, those participants indicated their pursuit of graduate education was motivated by hopes in better career opportunities. It is inconclusive that graduating the IBT_2017 enhanced participants’ employability, but the feedback collected 9 months upon the course end (filled by 30/51 of the successful participants) indicate that many of the then-students participants have completed their degrees and were offered jobs based on their new skills (Supplementary SF12, SF13). The same pattern was observed with the IBT_2016 MSc and PhD alumni who received teaching positions offers in local newly established faculties to teach bioinformatics-related courses or computational laboratories (personal communications).

### 5.2 Participants’ perceptions & expectations

By design, the IBT course is taught over 3 months, to train basic bionformatics users. Besides providing relevant and high standard content, the IBT core team and various instructors have placed great emphasis on defining clear learning objectives and outcomes prior to each taught module, and maintained the logical structure of the course despite removing the structural proteomics module from the IBT_2017 [Gurwitz *et al.*, 2017].

Therefore course participants had clear expectations out of each module, manifested as largely positive indicators in their responses to our surveys (Figure 3, and Supplementary SF6-SF11), and high success rate (excluding early withdrawals, 94% of the participants satisfied the course requirements, see also section 5.1). Considering that the majority of the participants were current graduate students (Figure 1C, Supplementary SF3) in a domain related to genetics or molecular biology (Supplementary SM8), they were exposed to concepts and uses of Databases; but less familiar with Linux, Genomics and related topics, because there is none or limited postgraduate degrees in bioinformatics currently in Sudan.

Regarding utilizing local and remote classroom resources, we make few observations. While we see that our participants were comfortable networking with each other, we see less interactions with participants from other classrooms. Partially, this is attributable to the demographics of the classrooms (section 5.1). While a few minutes were spared at the beginning of each session for the scattered classrooms to introduce themselves via Mconf, this was ineffective in linking our participants with other classrooms (Figure 4) because often times the connection would be too noisy for a meaningful conversation. The same issue often arose during the discussion time at the end of each session when the instructors are typically available and answering questions live. Alternatively, our participants tended to discuss issues through the forum or the chat rooms available through Vula. There is more engagement in these avenues towards the end of the course from Figure 4.

Another element explaining the high success rate of the 2 classrooms is the local help provided to the participants through the local TAs who were alumni of the 2016 iteration. From one perspective, testimonials from successful alumni have been shown to improve the sense of belonging in MOOCs, and there by learners’ performance [Kizilcec *et al.*, 2017]. This is especially true as the IBT_2017 included training sessions for the local team in each classroom on how to best facilitate the course. These training sessions employed the Mental Contrasting with Implementation Intentions (MCII) model [Kizilcec and Cohen, 2017] in equipping the local staff with best strategies for course facilitation by asking them to set goals for the course, and then predict future challenges and devise contingency plans. While not all identified challenges were within direct control (see section 5.3), the exercise gave a sense of confidence to the local staff. Additionally, the majority of our volunteering staff already had prior teaching experience (7/8 TAs, in addition to the class PI). The responses from those TAs indicate that they benefited from their previous experience, so they were comfortable facilitating demographically diverse classes (Supplementary SF14). On the personal level, the TAs were highly satisfied with the experience, even though they had other pressing commitments playing at the same time as the IBT_2017 (Supplementary SF15).

A final element in examining the IBT_2017 is its non-cognitive factors. These include the expected load incurred by participating in the IBT, which many found to be overwhelming when compared with commercial 1 week bioinformatics courses with no exams or assignments. This perception has exacerbated especially with the Linux module as can be seen in Figure 2. Another factor may be the course language. Sudan is officially an Arabic speaking country, though university educated graduates have some competency in English. However, one teaching style commonly employed, especially at the undergraduate level, is to deliver teaching curricula in Arabic, while maintaining English keywords and having English-based handouts (Supplementary SF16). This means that for many of the participants, complete course delivery in English is challenging. Collectively, these factors could explain the large withdrawal from the course before the mid-point (26% of the total participants, Figure 2). A possible workaround might be for the course participants to take a preliminary Scientific English course, either in a professional center in Sudan, or via MOOCs, like the positively reviewed Couresera offering of *“English for Science, Technology, Engineering, and Mathematics”* available on (https://www.coursera.org/learn/stem)

### 5.3 Reflections on local logistics

The IBT_2017 used a blended MOOC [Ghadiri *et al.*, 2013], or multi-delivery model [Gurwitz *et al.*, 2017], for learners to access, discuss and submit their assessments and tests. The online resources used: Mconf for example, were open source and were not network intensive; which made them appropriate to the local set up. Also, the geographical distribution of modules’ instructors across Africa provided a familiar context for participants to relate to. These aspects, technological infrastructure and relevant context, are effective in making a MOOC accessible to participants, and hence improving performance [Castillo *et al.*, 2015]. We didn’t compare with similar students’ performance on the offerings from edX and Coursera (table 1), but it is interesting to pursue in the future.

By large, one can see that the design and running of the IBT_2017 followed the *Ten simple rules for developing a short bioinformatics training course* of Via *et al.* [2011]. Particularly, we comment on the application of the following rules (as they pertain to the local classrooms in the University of Khartoum):

- **Setting Practical and Realistic Expectations achieved by specifying target audience (Rule1, Via *et al.* [2011]):** The target audience for the IBT are participants with molecular biology background. Those were the participants able to satisfy the course requirements, while graduates from certain faculties (like Computer Science) withdrew early on. Otherwise, in our case, it did help in yielding a high success rate that we had a long waiting list in place.
- **Class diversity:** in terms of academic backgrounds, graduating universities and career stage (1; Supplementary SF3). Additionally, proper advertising, and conscious selection of participants (in lights of the drop out factors we highlighted) should help in a better educational outcome.
- **Ensuring Computational Equipment Preparedness and Support Availability (Rule3, Via *et al.* [2011]):** Before and throughout the IBT,the local IBT_2017 team would meet weekly and update on resources needed and tasks assigned. A myriad of platforms were used for this: Trello boards for planning and follow up (https://trello.com/), a Google mailing list for emails, and Authorea (https://www.authorea.com) for collaboratively working on this manuscript. For some of the members, a chatting app like Whatsapp (https://www.whatsapp.com/) was needed, but overall, experiencing other platforms was deeply appreciated and highly regarded.
- **Allowing Interactivity and Providing Time for Reflection, Individual Analysis, and Exploration (Rule 9, Via *et al.* [2011]):** This was achieved via the in-person sessions, with some of the IBT_2016 alumni as teaching assistants. Those sessions assured accessible local support to the IBT_2017 participants and thereby reduced their anxiety which is otherwise a major MOOC withdrawal reason [Hew and Cheung, 2014a]. Also, for learners from less developed countries, success stories from previous course alumni have been shown to improve performance [Kizilcec and Cohen, 2017]
- **Career progress and capacity building:** The IBT aims to equip African researchers with the skills and knowledge to launch their careers and establish their science. The 30 responses from the follow up survey sent 9-months upon the end of the IBT_2017 show that many participants have moved from being students (53.3% at the time of the IBT) to being junior or middle staff (collectively 56.3%)(Supplementary SF12). For some of them, the IBT helped in securing new job offerings (Supplementary SF13).

## 6 Best practices

### 6.1 Online course design elements

#### Sessions and Modules learning outcomes

At the first session of each IBT module, trainers identified and emphasized on the learning outcomes [Gurwitz *et al.*, 2017]. This gave participants clear expectations, so they were able to evaluate their progress which is a good course success indicator. This was also reflected by the percentage of participants who were able to meet the course requirements.

#### Trainers interactive sessions with course participants

Trainers were available for live discussions with participants and could activate their cameras for more personal experience [Gurwitz *et al.*, 2017].

#### Hands-on sessions & Teaching assistants

During this free 3-months course, a large amount of time was dedicated to hands-on sessions, where participants practice what they are learning [Gurwitz *et al.*, 2017] with the help of local TAs. This is of great importance, as often trainees fail to appreciate the applicability of theoretical material to real data.

#### Video Conferencing System

The IBT classrooms connected trainers to all African classrooms via Mconf, an open-source video conferencing platform (https://mconf.sanren.ac.za/), and classrooms either activated their microphones or entered text into a chat box to ask questions [Gurwitz *et al.*, 2017]. Considering it was a free resource, Mconf features were mostly satisfactory (real-time chat, screen sharing, file sharing and classroom mode). Issues with sound clarity and disconnection is motivating the IBT core team to consider alternative platforms in future IBT iterations (personal communication).

#### Learning Management System

Throughout the IBT, Vula was utilized to send out announcements, manage participants, track progress, and allow interactions amongst participants, trainers and staff [Gurwitz *et al.*, 2017].

### 6.2 Local settings

#### Registration database

We had many people interested in the IBT. At the time of writing, there are about 400 interested applicant in our waiting list, coming from different universities, career and educational backgrounds.

#### Teaching assistants IBT alumni

During the planning of the IBT_2017 course, we came up with the idea of taking IBT_2016 alumni as volunteering TAs. This improved the experience of the IBT_2017 participants.

#### Local Logistics and Planning

Compared to the IBT_2016 course intake of only 22 participants, we were successful to intake 73 participants for the IBT_2017. This increase is credited to a partnership with University of Khartoum Main Library.

#### Class diversity

course participants with different expertise and research interests contribute to a more interactive classroom setting, as learners help each other, and build on each others’ complimentary skills.

## 7 Conclusion and Recommendations

### 7.1 Challenges

1. **Location:** Finding computer labs to accommodate a larger intake of the IBT participants is a challenge, because its duration is bound to overlap with parts of the academic year in most of the relevant faculties, and hence they can’t offer their labs for the entire 3-months duration.
2. **Timing:** Some of the course participants are full time MSc students, and some of the IBT sessions collided with the timing of their exams (~25 participant). The flexibility and sensitivity of the IBT core team in giving them some grace period helped these participants make up for missed activities.

### 7.2 Lessons learnt

1. Working closely with concerned entities within the University provides support in allocating more infrastructure and resources.
2. More active collaborations with governmental entities like the Ministry of Higher Education sustains training and research efforts like the IBT.

### 7.3 Looking forward

1. Collaborations with other universities/ Research centers (in other states besides Khartoum).
2. Arranging similar courses (multi-delivery model) in other areas like data science and health informatics.

## 8 Key points

- Learners in less developed countries are keen on to seizing educational opportunities. For MOOCs to be attractive, some effort is needed.
- Blended learning employing multi-delivery models can be effective in bridging the achievement gap in MOOCs, even in bioinformatics courses.
- The training of local TAs from course alumni improves the sense of belonging to a MOOC, and hence improves learners’ performance.
- Working in tandem with other local bodies to secure the needed infrastructural resources is another factor contributing to a successful training.

## Author contributions

- Conceptualization: AA, FMF
- Formal analysis: AA, FMF
- Funding acquisition: FMF
- Investigation: AA, AAA, MTE, MAS, AEK, HME, BDH, FMF
- Methodology: AA, AAA, MTE, MAS, AEK, HME, BDH, FMF
- Project administration: FMF
- Resources: FMF
- Software: AA
- Supervision: FMF
- Writing/ Original draft: AA, AAA, FMF, MAS, MTE, HME
- Writing/ Review and editing: AA, AAA, MTE, MAS, AEK, HME, BDH, FMF

## Acknowledgements

- IBT_2017 core team: Kim T. Gurwitz (University of Cape Town), Shaun Aron(University of Witwatersrand), Sumir Panji (University of Cape Town), Suresh Maslamoney (University of Cape Town), Nicola Mulder(University of Cape Town)
- IBT 2017 trainers
- IBT 2017 Sudan Course participants
- University of Khartoum Faculty of Science
- University of Khartoum Main library
- University of Khartoum Information Technology Network Administration (ITNA)
- Dr. Pamela Greenwell, University of Westminister (UK) for her insightful comments and review of an initial version of the manuscript

